# Morphogen-driven melanin pathway dynamics regulated by *Wnt1* and *apontic-like* underlie larval spot coloration in *Bombyx mori*

**DOI:** 10.1101/2025.04.03.645855

**Authors:** Shinichi Yoda

## Abstract

In insects, conspicuous larval pigmentation patterns serve critical ecological roles such as warning signals and mimicry, yet their underlying genetic regulation remains poorly understood. In this study, I investigated the molecular mechanisms underlying black and yellow pigmentation patterns in three distinct larval spot types of the silkworm *Bombyx mori*: large, diffuse *L*-spots of the *Multilunar* (*L*) mutant; small, sharply defined +*^p^*-spots of the *Normal* strain; oval *p^M^*-hybrid spots of an interspecific hybrid with *Bombyx mandarina*. Each spot type comprises a yellowish center surrounded by a black periphery, forming crescent-shaped pigmentation patterns. Chemical treatments confirmed that both colors are melanin-based. Using quantitative PCR and RNA interference (RNAi), I analyzed six melanin synthesis genes (*Tyrosine Hydroxylase*, *Dopa Decarboxylase*, *laccase2*, *yellow*, *tan*, and *ebony*) and discovered that black pigmentation involves both dopa/dopamine- and NBAD-melanin synthesis, whereas yellow pigmentation primarily reflects only the latter. I further examined *Wnt1* and *apontic-like* (*apt-like*) using qPCR, RNAi, and TALEN-mediated mosaic analysis. *Wnt1* expression localized to presumptive spot areas, functioning dose-dependently to regulate both spot size and pigment composition: high *Wnt1* levels induced larger spots with yellow centers, while reduced *Wnt1* expression resulted in black pigmentation and smaller spots. *Wnt1*-activated transcription factor *apt-like* was required for pigmentation in all spot types without influencing spot size. Taken together, the results of this study reveal a morphogen-driven gene regulatory network in which *Wnt1* dosage and downstream transcriptional cascades orchestrate pigment placement and patterning, offering new insights into the modular genetic control of insect pigmentation.

## 1. Introduction

Color pattern formation in insects is pivotal for diverse ecological functions, including predator avoidance, thermoregulation, and intraspecific signaling (Postema et al., 2022). Among these, conspicuous pigmentation patterns in caterpillars deserve particular attention for their role in defense, functioning as warning signals, disruptive camouflage, or mimicry-based deterrents (Ruxton et al., 2019). Aposematic coloration typically involves high-contrast combinations (e.g., red, yellow, white, and black), enhancing visibility against natural backgrounds and reducing recognition errors by predators (Stevens and Ruxton, 2012; Ruxton et al., 2019). The association between bright coloration and defense has been recognized since the early observations of Wallace, linking conspicuous pigmentation to unpalatability in nonreproductive caterpillars (Caro and Ruxton, 2019). The widespread presence of such traits across taxa suggests convergent evolution, with similar selective pressures driving the independent emergence of bright coloration in multiple lineages (Poulton, 1890; Cott, 1940; Edmunds, 1974; Stevens and Ruxton, 2012; Aronsson and Gamberale-Stille, 2013; Arenas et al., 2014).

In lepidopteran larvae, conspicuous markings (e.g., eyespots) often serve as predator deterrents by mimicking vertebrate eyes, eliciting predator hesitation or avoidance (Janzen et al., 2010; Stevens and Ruxton, 2012). For instance, certain swallowtail butterfly caterpillars develop eyespots that resemble snake eyes, while others display bold red or orange spots that might mimic toxic species (Prudic et al., 2007). These striking markings are sometimes accompanied by striped patterns that appear cryptic from a distance but become highly conspicuous at close range, thereby enhancing their warning effect (Rothschild, 1975; Barnett and Cuthill, 2014).

While the ecological significance of these visual traits is well documented, the molecular mechanisms underlying the formation of high-contrast pigmentation patterns in caterpillars remain poorly understood. Specifically, how genetic factors regulate spot size and pigmentation composition (e.g., distinguishing black and yellow pigments) remains elusive.

Due to its well-characterized genome and numerous available pigmentation mutants, the silkworm (*Bombyx mori*) is a powerful model for studying the genetic mechanisms of larval color pattern formation (Goldsmith and Wilkins, 2006; International Silkworm Genome Consortium, 2008; Kawamoto et al., 2019). To date, over ten genes have been identified at larval pigmentation-involved loci (e.g., Futahashi et al., 2008, 2010; Liu et al., 2010; Yuasa et al., 2016; KonDo et al., 2017; Wu et al., 2024). Among these, the dominant mutant *Multilunar* (*L*), located at the *L* locus on chromosome 4 (Tazima, 1964), develops large dorsal spots (*L-* spots) across multiple body segments (Fig. 1, *L*). Each *L-*spot features an inner yellowish-brown region (hereafter referred to as yellow region), surrounded by an outer black region. The yellow-black color boundary is diffuse and the black region does not form a complete ring around the center (Fig. 1, *L*-spots). In contrast, the *Normal* (+*^p^*) phenotype, associated with the *p* locus on chromosome 2 (Tazima, 1964), displays smaller and more sharply defined spots limited to specific segments: eye spots, crescents, and star spots on the second thoracic, second abdominal, and fifth abdominal segments, respectively (Fig. 1, +*^p^*). Similar to *L-*spots, +*^p^-*spots comprise a yellow center and a black periphery, although their boundaries are clearly demarcated (Fig. 1, +*^p^*-spots). Beyond these two well-characterized patterns, a *B. mori* × *B. mandarina* hybrid strain, carrying the *p^M^* (*Moricaud*) allele at the *p* locus (Tazima, 1964), exhibits a brown ground color and spot patterns on the same segments as +*^p^* (Fig. 1, Hybrid). While *p^M^*-hybrid spots also feature a yellow center and black periphery, their shapes tend to resemble crescent-formed ovals rather than the sharply defined crescents of the +*^p^* strain (Fig. 1, *p^M^*-hybrid spots).

**Fig. 1.**
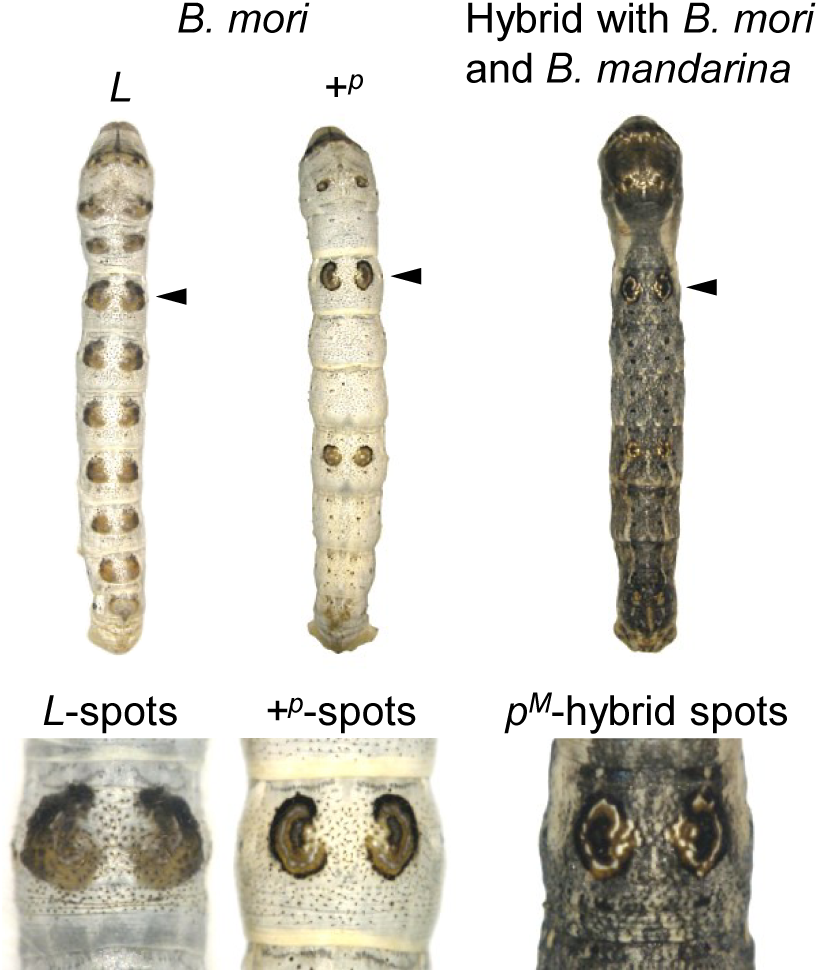
Three characteristic larval spot marking types in silkworms. From left to right: dorsal views of fifth instar larvae from the *Multilunar* (*L*) mutant at the *L* locus, the *Normal* (+*^p^*) strain at the *p* locus, and a hybrid strain between *Bombyx mori* and wild *B. mandarina*, carrying the *Moricaud* (*p^M^*) allele at the *p* locus. The *L* mutant exhibits multiple spots across consecutive thoracic and abdominal segments, whereas +*^p^* and the *p^M^*-hybrid display spots only on the second thoracic as well as the second and fifth abdominal segments. The magnified views below each larva present crescent-shaped spots (indicated with arrowheads) on the second abdominal segment. All spot types consist of an inner yellow region surrounded by an outer black region, partially in the *L*- and almost completely in the +*^p^*- and *p^M^*-hybrid spots.

*Wnt1* (*wingless*) is reportedly a key gene responsible for periodic *L-*spot formation, where its localized upregulation, induced by the molting hormone ecdysteroid, is restricted to the epidermis of presumptive spot areas (Yamaguchi et al., 2013). Wnt signaling is central to insect color pattern formation. In butterflies, *Wnt1* regulates wing eyespot formation and wing pattern variation by defining spatial pigmentation gradients (Monteiro et al, 2006; Martin and Reed, 2014; Özsu et al., 2017; Iijima et al., 2018). In *Drosophila guttifera*, Wnt signaling directs pigmentation patterning on the dorsal thorax and wings (Werner et al., 2010; Koshikawa et al., 2015). In addition, in nymphalid butterflies including *Heliconius*, *WntA* reportedly controls color pattern specification (Martin et al., 2012; Gallant et al., 2014; Mazo-Vargas et al., 2017).

Similarly, *apontic-like* (*apt-like*) is reportedly the key gene responsible for the *p* locus, regulating +*^p^* and *p^M^* spot pattern formation in *B. mori* (Yoda et al., 2014). *apt-like* encodes a transcription factor and functions as a pigmentation regulator. The *p* locus contains multiple alleles with a clear dominance hierarchy: *p^M^* is dominant over +*^p^*, which in turn is dominant over the recessive *p* allele. The genetic interaction between *Wnt1* and *apt-like* is proposedly essential for +*^p^-*spot formation, where *Wnt1* expression in the presumptive +*^p^-*spot area induces *apt-like* expression (Yoda et al., 2014), ultimately leading to sharply defined spot development in specific body segments.

In this study, I investigated the genetic basis of black and yellow pigmentation in *B. mori*. I identified key pigments and related genes through chemical treatment, qPCR, and RNAi analysis, focusing on *L-* and +*^p^-*spots. Furthermore, I examined the roles of *Wnt1* and *apt-like* in regulating spot size and pigmentation across three spot types. By quantifying *Wnt1* and *apt-like* expression and performing RNAi knockdowns, I evaluated their contributions to spatial and color pattern differentiation.

## 2. Materials and Methods

### 2.1. Insects

The following silkworm strains were obtained from the Institute of Genetic Resources at Kyushu University: the *Multilunar* (*L*) mutant strain g01 (*L*/*L*, +^p^/+*^p^*), the *Normal* (+*^p^*) strains u41 and t31 (+*^L^*/+*^L^*, +*^p^*/+*^p^*), and the *p^M^* introgression strain T02 (+*^L^*/+*^L^*, *p^M^*/+*^p^*) (Fujii et al., 2021). In addition, an interspecific F1 hybrid was generated by crossing a *B. mandarina* female (field-collected in Kashiwa City, Japan) with a *B. mori* N4 strain male (+*^L^*/+*^L^*, *p*/*p*).

### 2.2. Chemical treatments for estimating pigment types in *L*- and +*^p^*-spots

To separate the epidermal tissues from the cuticle, fifth instar larval integuments were soaked in 10% sodium dodecyl sulfate (SDS) at room temperature for at least 12 h. The epidermal layer was then carefully removed using fine tweezers. To determine whether the pigments remaining in the cuticle were melanin-based, the cuticle was hydrolyzed in 1 M hydrochloric acid (HCl) at 90 °C for 45 min, following the protocol of Koch and Kaufmann (1995). The treated cuticular specimens were examined using a KEYENCE VH-5500 microscope system (KEYENCE, Japan).

### 2.3. RNA isolation and quantitative reverse-transcription polymerase chain reaction (qRT-PCR)

For RNA sampling, I used the *L* strain g01 at two feeding stages (C1 and C2) and three molting stages (D3, E1, and E2), as defined by the spiracle index (Kiguchi and Agui, 1981). The spiracle index provides a morphological time marker to infer precise developmental stages during the larval molt based on visible changes in the spiracle. Epidermal samples from the white, black, and yellow regions of the *L*-spots were dissected from the second and third abdominal segments and washed in cold phosphate-buffered saline (PBS). Total RNA was extracted using precooled TRI reagent (400 μL per sample; Sigma-Aldrich), treated with DNase I (0.2 U; Takara, Japan) at 37 °C for 15 min, and purified by phenol–chloroform extraction. Reverse transcription was performed using a random hexamer primer (N6) and the PrimeScript IV 1st Strand cDNA Synthesis Kit (Takara, Japan), with the following thermal profile: 30 °C for 10 min, 42 °C for 20 min, and 70 °C for 15 min to inactivate the reaction. Quantitative PCR was conducted on a QuantStudio 5 System (Applied Biosystems) using PowerTrack SYBR Green Master Mix (Thermo Fisher Scientific). Cycling conditions were: 95 °C for 2 min, followed by 40 cycles of 95 °C for 5 sec and 60 °C for 30 sec. Data analysis was performed using the QuantStudio Design and Analysis Software (v1.5.1) with the relative standard curve method. Primer sequences are listed in Table S1. All qPCR data are presented as mean ± standard deviation (SD) from four biological replicates. Statistical analysis was performed using Dunnett’s test (two-sided), with *P* < 0.05 considered statistically significant.

### 2.4. *in vivo* electroporation-mediated RNA interference (RNAi)

Small interfering RNAs (siRNAs) were designed according to the criteria described by Yamaguchi et al. (2011). Target genes included *Tyrosine Hydroxylase (TH)*, *Dopa Decarboxylase (DDC)*, *laccase2*, *yellow*, *tan*, *ebony*, *Wnt1*, and *apt-like* (see Table S2 for sequences). A universal negative control siRNA (Nippon Gene, Japan) was used as a negative control. Each siRNA (250 μM, 0.75–1.0 μL) was injected into the hemolymph from the lateral side of the fifth abdominal segment in fourth instar larvae, immediately after the third molt, using a glass needle connected to a microinjector (FemtoJet, Eppendorf).

Immediately following injection, a droplet of PBS was applied over the spot markings on the second abdominal segment. Electroporation was then performed using the *in vivo* somatic transgenesis method described by Ando and Fujiwara (2013), with the positive electrode placed on the left-side spot. Electroporation parameters were as follows: 20 V for *L* and hybrid strains, 18 V for +*^p^*strains, 280 ms pulse length, 1 min interval, and 5 pulses in total. Phenotypic effects were assessed in fifth instar larvae using a VH-5500 microscope system (KEYENCE, Japan).

### 2.5. Measurement of Spot Area in Wnt1 RNAi Individuals

To quantify changes in spot size following *Wnt1* RNAi, the areas of *L*-spots, +*^p^*-spots, and *p^M^*-hybrid spots were measured using Fiji (ImageJ). The spot on the untreated (control) side of each larva was used as an internal reference for comparison with the RNAi-treated side. For imaging, the second abdominal segment, where spots are located, was photographed with the dorsal midline centered to enable symmetrical visualization of both sides. Spot areas were manually measured using the Freehand Selection tool in Fiji by carefully tracing the spot boundaries. Automated segmentation was avoided due to its poor accuracy in distinguishing spot edges. To reduce variability in manual tracing, each measurement was repeated three times per spot type, and the average value was recorded as the final area. Relative spot size was calculated as the ratio of the RNAi-treated spot area to the corresponding control spot area. This ratio was used to evaluate the extent of spot size reduction following *Wnt1* knockdown. All data are presented as box plots based on six (*L*-spots), four (+*^p^*-spots), and three (*p^M^*-hybrid spots) biological replicates. Statistical significance was evaluated using a two-sided paired *t*-test, with *P* < 0.05 considered statistically significant.

### 2.6. Transcription activator-like effector nuclease (TALEN)-mediated mosaic analysis

A TALEN pair targeting *apt-like* was designed using TALEN Targeter 2.0 (https://TALE-nt.cac.cornell.edu/). Potential off-target sites in the *B. mori* genome were assessed using BLAST, and both the left and right TALEN sequences were confirmed to lack significant homology elsewhere in the genome. TALEN mRNAs were synthesized and microinjected into embryos of the *p^M^* introgression strain (T02) following the protocol described by Takasu et al. (2013). For mutagenesis, TALEN left and right mRNAs (each at 200 ng/μL) were co-injected as a mixture. A total of 61 *p^M^*embryos were injected, of which three larvae hatched. One of the hatched individuals exhibited a mosaic pigmentation phenotype. Genomic DNA was extracted from this mosaic larva to assess TALEN-induced mutations at the target locus. The region surrounding the TALEN target site was amplified by PCR using the primers Bm3234_Exon6-F1 (5′-GACTATCGACAACGAACGACAGC-3′) and Bm3234_Exon6-R1 (5′-CGTTGTCGATAGTCGTCCGTTTG-3′). The PCR products were subcloned into the pGEM-T Easy vector (Promega) and sequenced. Eight clones were analyzed, and three of them (37.5%) carried deletions consistent with TALEN-induced mutations.

## 3. Results

### 3.1 Estimation of black and yellow pigment composition in *L-* and +*^p^-*spots

Previous research suggested that the *L- and +^p^-*spots contain both melanin and ommochrome pigments (Yoshitake et al., 1952; Ding et al., 2019). To determine whether the black and yellow pigments in the spot markings comprise melanin, ommochrome, or a combination of both, I dissected the integuments from the second abdominal segment of *L* and *+^p^* larvae (Fig. S1A). I treated the integuments with SDS to remove the epidermis, leaving only the cuticle. The black and yellow pigmentation remained visible in the cuticle with minimal color change after SDS treatment (Fig. S1B) compared to the untreated samples (Fig. S1A).

Furthermore, HCl hydrolysis of the SDS-treated cuticles did not elute either black or yellow pigments (Fig. S1C). Since melanin is a highly polymerized and HCl-insoluble pigment (Lea, 1945; Koch and Kaufmann, 1995), this result suggests that both black and yellow pigments in *L* and *+^p^* are melanin-based.

### 3.2. Temporal expression profiles of melanin synthesis genes in the black and yellow coloration of the *L-*spots

To decipher the molecular mechanisms underlying color variation between black and yellow pigmentation in the *L-*spots, I analyzed the expression of six key melanin synthesis genes: *Tyrosine Hydroxylase* (*TH*), *Dopa Decarboxylase* (*DDC*), *yellow*, *laccase2* (also known as *MCO2*), *tan*, and *ebony* (Fig. 2A). In insects, three representative melanin pigment types (i.e., dopa-, dopamine-, and N-β-alanyldopamine (NBAD)-melanins) are synthesized through distinct but interconnected enzymatic steps. The pathway begins with *TH* converting tyrosine into dihydroxyphenylalanine (DOPA), which is then converted into dopamine by *DDC*. DOPA gives rise to dopa-melanin, typically associated with dark brown coloration, while dopamine is the dopamine-melanin precursor, contributing to black pigmentation and is promoted by *yellow*. Alternatively, dopamine could be converted into NBAD by *ebony*, leading to NBAD-melanin formation, which is a lighter pigment. *tan* could reverse this reaction, regenerating dopamine from NBAD. Finally, *laccase2* catalyzes DOPA, dopamine, and NBAD oxidation into quinones that polymerize into their respective melanin pigments in the cuticle (Shamim et al., 2014; Futahashi et al., 2022).

**Fig. 2.**
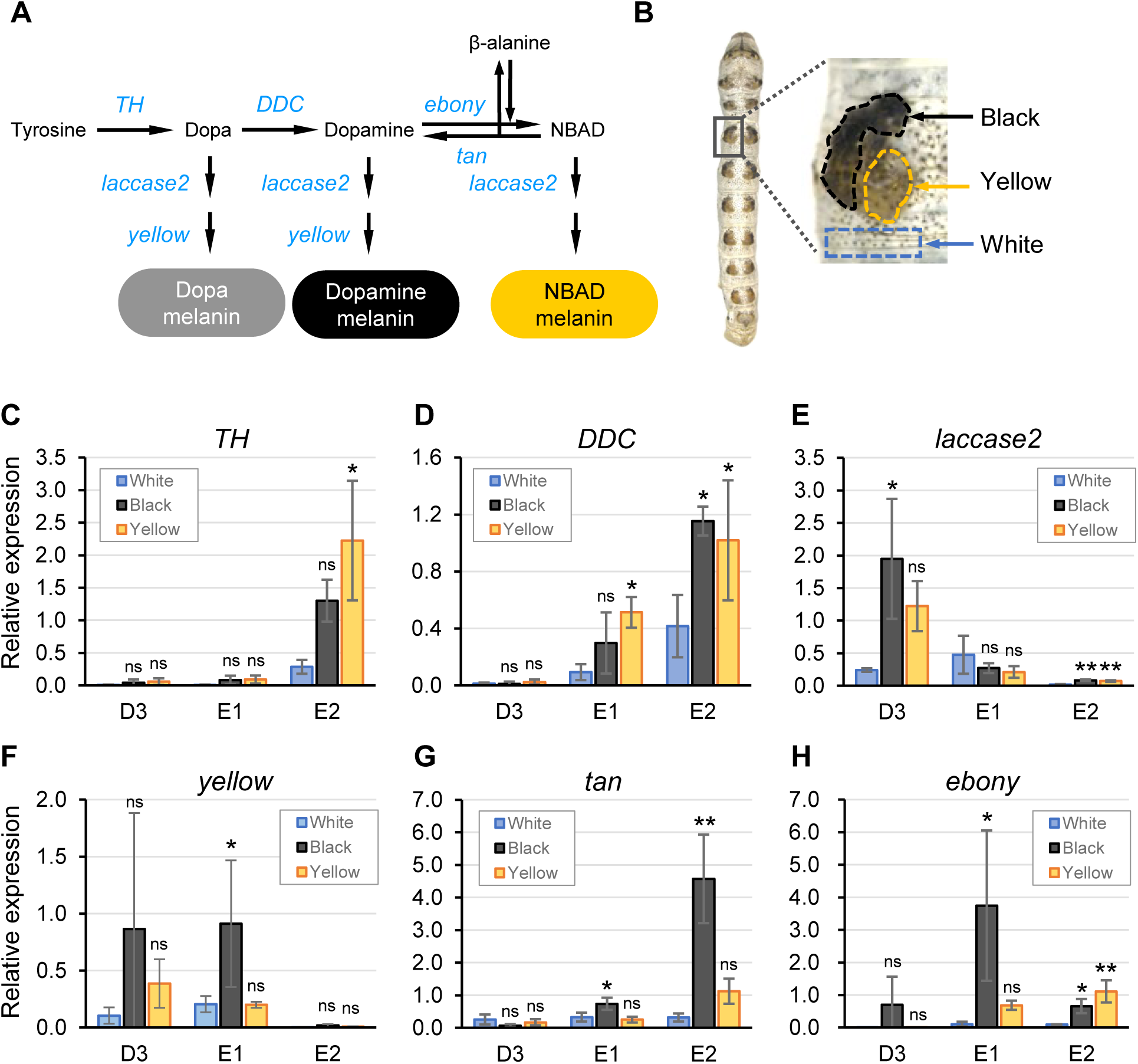
The differential expression of six core melanin synthesis genes correlates with the black and yellow regions in the *L*-spots. (**A**) Schematic representation of the insect melanin synthesis pathway, highlighting the six core melanin synthesis genes analyzed in this study. (**B**) Diagram presenting the dissected regions (i.e., black, yellow, and white) on the second abdominal segment of *Multilunar* (*L*) larvae at the fourth instar stage. (**C–H**) Relative expression levels of *Tyrosine Hydroxylase* (*TH*; **C**), *Dopa Decarboxylase* (*DDC*; **D**), *laccase2* (**E**), *yellow* (**F**), *tan* (**G**), and *ebony* (**H**) in the black and yellow regions, normalized to the white region, which served as the control. *RpL3* was used as the internal reference gene. Gene expression was measured at three developmental stages during the molting period (i.e., D3, E1, and E2), defined based on the spiracle index. The data are represented as the mean ± SD (n = 4). The error bars represent the standard deviation. Statistical significance was assessed using Dunnett’s test: *P* < 0.05; *P* < 0.01; ns, not significant.

I assessed gene expression in the black and yellow regions of the second abdominal segment in *L*, with the white region serving as the control (Fig. 2B). I used qPCR to quantify the expression levels during three molting stages: D3, the early stage before pigmentation begins; E1, pigmentation onset; E2, final cuticle melanization phase before molting (Kiguchi and Agui, 1981).

Both *TH* and *DDC* were upregulated at the late E2 stage, with *TH* strongly expressed both in the black and yellow regions, particularly in the latter (Fig. 2C). *DDC* expression increased earlier in the yellow region at E1 and continued to rise in both pigmented regions at E2 (Fig. 2D). *laccase2* displayed high expression both in the black and yellow region at D3, although its expression level varied in the latter (Fig. 2E).

In the black region, *yellow* and *tan* were significantly upregulated. *yellow* expression increased from D3 to E1 (Fig. 2F), while *tan* peaked between E1 and E2 (Fig. 2G). *ebony* was highly expressed in the yellow region at E2 and also upregulated in the black region from E1 to E2 (Fig. 2H). These patterns suggest that *ebony* contributes to NBAD-melanin synthesis both in the black and yellow regions. In addition, *tan* and *yellow* upregulation in the black region likely facilitates NBAD-melanin mixing with dopa- and dopamine-melanin, thereby contributing to the final pigmentation pattern observed in the *L-*spots.

In summary, these results represent a strong correlation between melanin synthesis gene expression and pigmentation in the *L-*spots, highlighting the interplay between NBAD-, dopa-, and dopamine-melanins in determining spot coloration.

### 3.3. RNAi knockdown effect of six melanin synthesis genes on the *L-* and +*^p^-*spots

To investigate the role of melanin synthesis genes in black and yellow pigmentation in the *L-* and +*^p^-*spots, I knocked down six core genes using RNAi. I performed *in vivo* electroporation-mediated siRNA delivery in the left spot region of the second abdominal segment of *L* and +*^p^* larvae within 24 h after molting to the fourth instar, and evaluated pigmentation at the fifth instar. I introduced a negative control siRNA to exclude nonspecific RNAi effects on pigmentation in the *L-* and +*^p^-*spots (Fig. S2).

*TH* knockdown significantly suppressed black pigmentation both in the *L-*spots and +*^p^-*spots, whereas it relatively moderately affected yellow pigmentation (Fig. 3A). These results indicate that *TH* is essential for black pigmentation and also contributes to yellow pigmentation. The limited impact on yellow regions might be due to the high *TH* expression not only during the E2 stage but also immediately after molting, suggesting that *TH* might have contributed to yellow pigment formation before a pronounced RNAi effect onset.

**Fig. 3.**
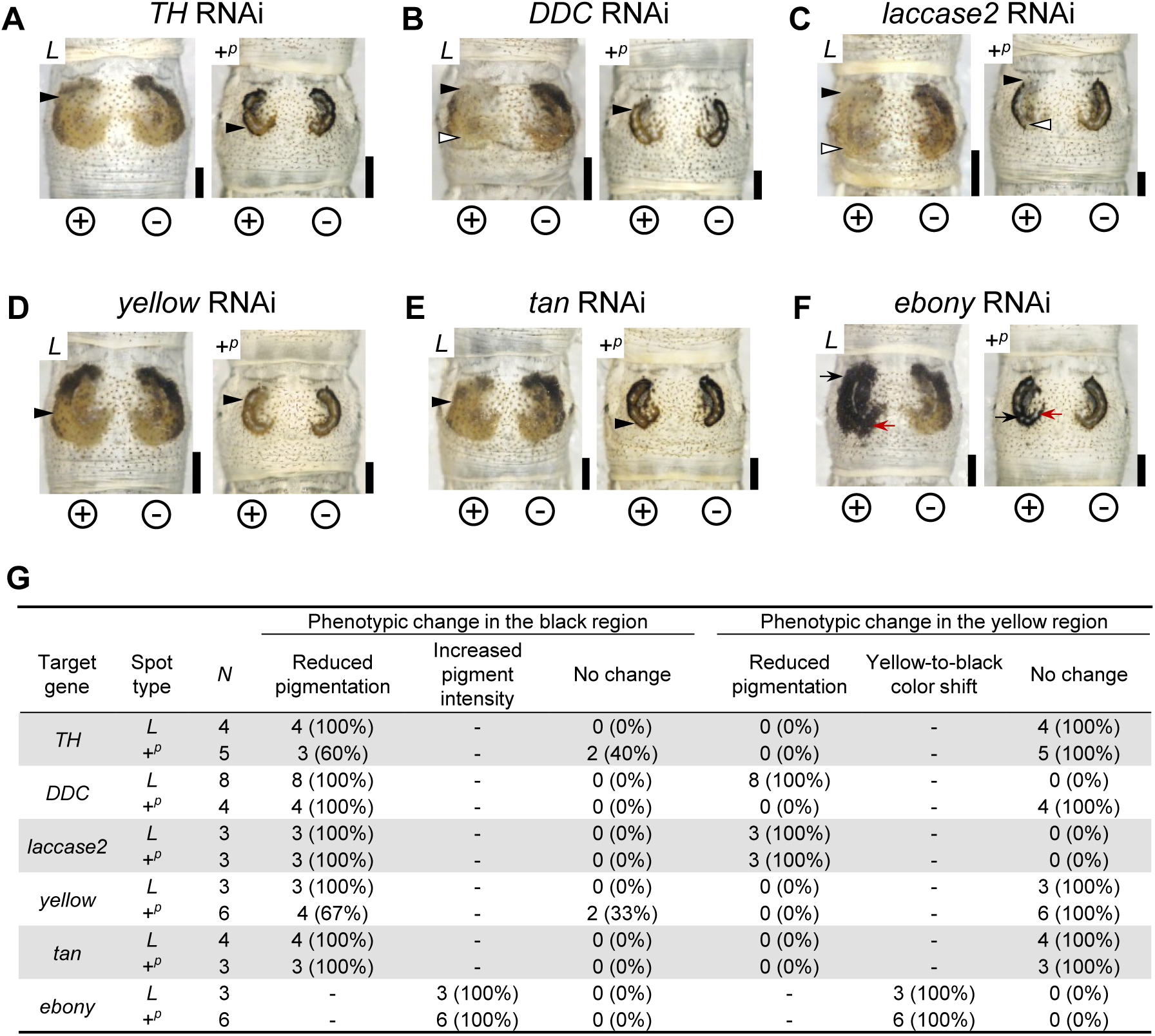
RNAi knockdown effect of six melanin synthesis genes on the *L-* and +*^p^*-spots. (**A–F**) Representative phenotypes upon RNAi-mediated knockdown of six melanin synthesis genes: *TH* (**A**), *DDC* (**B**), *laccase2* (**C**), *yellow* (**D**), *tan* (**E**), and *ebony* (**F**) in *L* and +*^p^* larvae. RNAi was performed via *in vivo* electroporation by delivering siRNA into the left-side spots; ‘+’ and ‘–’ indicate the direction of the applied electric field during electroporation. Black and white arrowheads indicate reduced pigmentation in the black and yellow regions, respectively. Black and red arrows indicate increased black pigmentation and a color shift from yellow to black, respectively. Scale bars: 1 mm. (**G**) Summary of RNAi-induced phenotypic changes in the black and yellow regions of the *L*- and +*^p^*-spots.

*DDC* knockdown in the *L*-spots suppressed both black and yellow pigmentation (Fig. 3B, *L-* spots), consistent with its high expression levels in these regions. This result indicates the necessity of *DDC* for the formation of both pigment types in the *L-*spots. In the +*^p^*-spots, *DDC* knockdown primarily reduced black pigmentation, less pronouncedly affecting the yellow region (Fig. 3B, +*^p^*-spots). This observation implies that while *DDC* contributes to black pigmentation in +*^p^*, its role in yellow pigmentation might be partially compensated or subject to different regulatory mechanisms. These differences might reflect variations in *DDC* expression timing between the *L* and +*^p^* alleles.

*laccase2* knockdown in the *L*-spots severely reduced both black and yellow pigmentation, consistent with its high expression in these regions (Fig. 3C, *L*-spots). I observed a similar effect in the +*^p^*-spots, where *laccase2* knockdown also reduced both pigment types (Fig. 3C, +*^p^*-spots). These results confirm that both black and yellow pigmentation in the *L*- and +*^p^*- spots are melanin-based.

*yellow* and *tan* knockdown in the *L*-spots specifically inhibited black pigmentation without affecting yellow pigmentation, consistent with their high expression in the black regions (Figs. 3D and E, *L*-spots). In the +*^p^*-spots, *yellow* and *tan* knockdown similarly suppressed black pigmentation (Figs. 3D and E, +*^p^*-spots), further supporting their role in black pigment formation.

*ebony* knockdown yielded contrasting results. In the *L*-spots, it induced a loss of yellow coloration simultaneously with increased black pigmentation both in the black and yellow regions (Fig. 3F, *L*-spots). I observed a similar pattern in the +*^p^*-spots, where *ebony* knockdown enhanced black pigmentation across both regions (Fig. 3F, +*^p^*-spots). These findings indicate that *ebony* suppresses black pigmentation in the yellow regions by promoting NBAD-melanin synthesis, thereby preventing excessive black pigment accumulation.

Taken together with the qPCR results, these findings suggest that *TH*, *DDC*, *laccase2*, *yellow*, *tan*, and *ebony* are crucial for black, while *TH*, *DDC*, *laccase2*, and *ebony* also regulate yellow pigmentation. These results further indicate that yellow pigmentation is primarily involved in NBAD-melanin, whereas black consists of both dopa/dopamine- and NBAD-melanins.

Figure 3G summarizes the phenotypic effects of RNAi knockdowns across all six melanin synthesis genes (including the frequency of pigmentation loss, intensity changes, and unaffected cases), providing a contribution overview of each gene to black and yellow pigmentation, thereby reinforcing their roles in melanin synthesis. While the RNAi knockdown of these genes affected pigmentation intensity and distribution, it did not alter spot size either in *L* or +*^p^* (Fig. 3G), suggesting that melanin synthesis genes regulate pigment production but not spatial patterning, and that melanin synthesis-independent upstream patterning mechanisms determine spot size.

### 3.4. *Wnt1* and *apt-like* expression patterns correlate with the black and yellow coloration in the *L*-spots

Ecdysteroid-regulated periodic *Wnt1* expression is reportedly responsible for generating twin spots (*L-*spots) in the silkworm *L* mutant (Yamaguchi et al., 2013). In addition, the gene *apt-like*, encoding a putative transcription factor and being responsible for the *p* locus, is pivotal for regulating pigmentation patterns, including +*^p^*-spot formation (Yoda et al., 2014). Notably, *L-*spots appear only when the +*^p^* allele is present (either in heterozygous or homozygous forms), but are absent in individuals carrying homozygous *p* alleles (Fig. S3), suggesting that *apt-like* in the +*^p^* genetic background is essential for *L-*spot formation. However, these previous studies did not functionally verify, through RNAi experiments, whether *apt-like* expression is necessary for *L-*spot formation or *Wnt1* expression is indeed required for +*^p^*-spot formation. Furthermore, the expression levels of *Wnt1* and *apt-like* in the black and yellow regions of the *L*-spots had not been characterized.

To address this, I used qPCR to analyze the *Wnt1* and *apt-like* expression patterns in the black and yellow regions of the *L-*spot during different developmental phases: the feeding stages (C1 and C2), when these genes are reportedly highly expressed, and the molting phase (D3), when their expression levels decline (Yamaguchi et al., 2013; Yoda et al., 2014). I compared the gene expression levels in the black and yellow regions of the second abdominal segment to those in the corresponding white region.

The results indicated increased *Wnt1* and *apt-like* expression levels both in the black and yellow regions compared to the white. In the yellow region, *Wnt1* expression increased from C1 to C2, then declined by D3, displaying a temporal peak (Fig. 4A). In the black region,

**Fig. 4.**
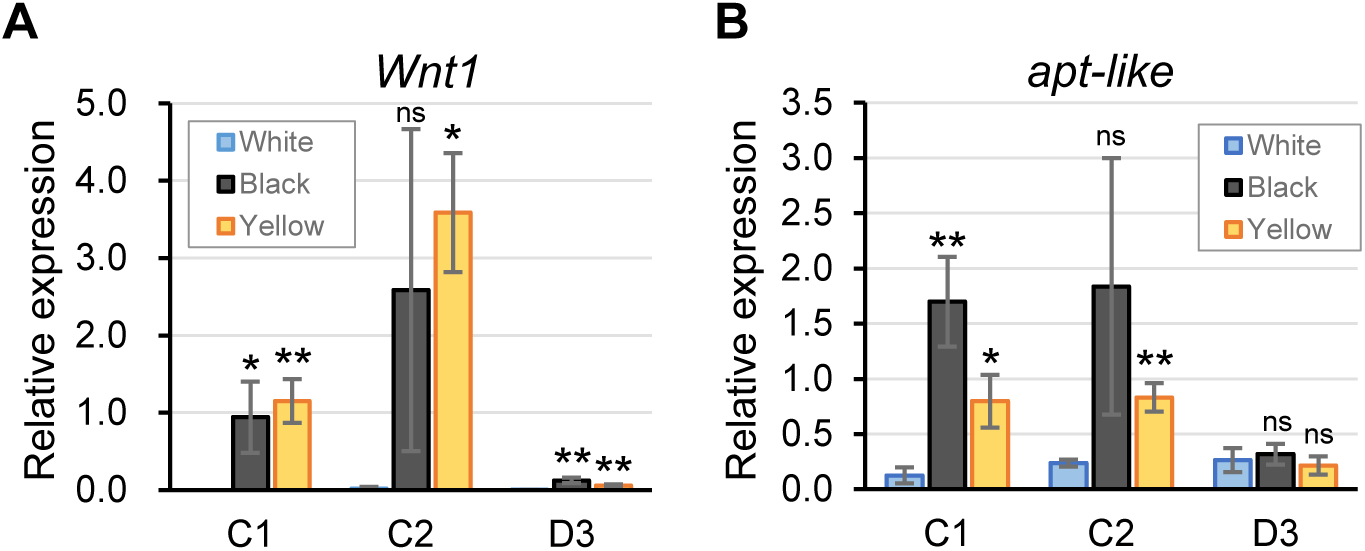
*Wnt1* and *apt-like* expression profiles in distinct *L*-spot pigmentation regions. (**A and B**) Relative expression levels of *Wnt1* (**A**) and *apt-like* (**B**) during the C1, C2, and D3 stages of the fourth instar in the *L*-spots. The epidermal samples were dissected as described in Fig. 2B. Both *Wnt1* and *apt-like* exhibited increased expression in the yellow region from C1 to C2. The expression in the black region also remained high across these stages, although the increase at C2 was not statistically significant. *RpL3* was used as the internal reference gene. The data are represented as the mean ± SD (n = 4). The error bars represent the standard deviation. Statistical significance was assessed using Dunnett’s test: *P* < 0.05; *P* < 0.01; ns, not significant.

*Wnt1* expression remained high across the stages, although the increase at C2 was not statistically significant. Similarly, *apt-like* was highly expressed both in the yellow and black regions during C1 and C2, followed by a decline at D3 (Fig. 4B). Notably, *Wnt1* tended to be more highly expressed in the yellow than in the black region, whereas *apt-like* exhibited the opposite trend, with higher expression in the black region than in the yellow.

### 3.5. Differential *Wnt1* and *apt-like* roles in spot pigmentation and size regulation

Next, as *Wnt1* and *apt-like* were highly expressed both in the black and yellow regions during critical developmental stages, I sought to functionally validate their pigmentation-related roles through gene knockdown experiments. *Wnt1* expression quantities in *L* are reportedly significantly higher than those in *+^p^*, suggesting a potential dosage-dependent effect on pigmentation. To examine how *Wnt1* could affect black and yellow pigmentation, I applied *Wnt1* siRNA to the left *L-*spot on the second abdominal segment during the fourth instar using *in vitro* electroporation and observed its impact in the fifth instar. Compared to the untreated side, the *L-*spot size decreased and its pigmentation shifted from yellow to black (Fig. 5A and B, *L-*spots). Similarly, *Wnt1* knockdown in the *+^p^-*spots yielded a reduced size, though with a less pronounced effect (Fig. 5A and B, +*^p^-*spots). Furthermore, in a hybrid strain crossed with *Bombyx mandarina*, naturally exhibiting larger spots than *+^p^*, *Wnt1* knockdown caused a slight overall reduction in spot size (Fig. 5A and B, *p^M^*-hybrid spots).

**Fig. 5.**
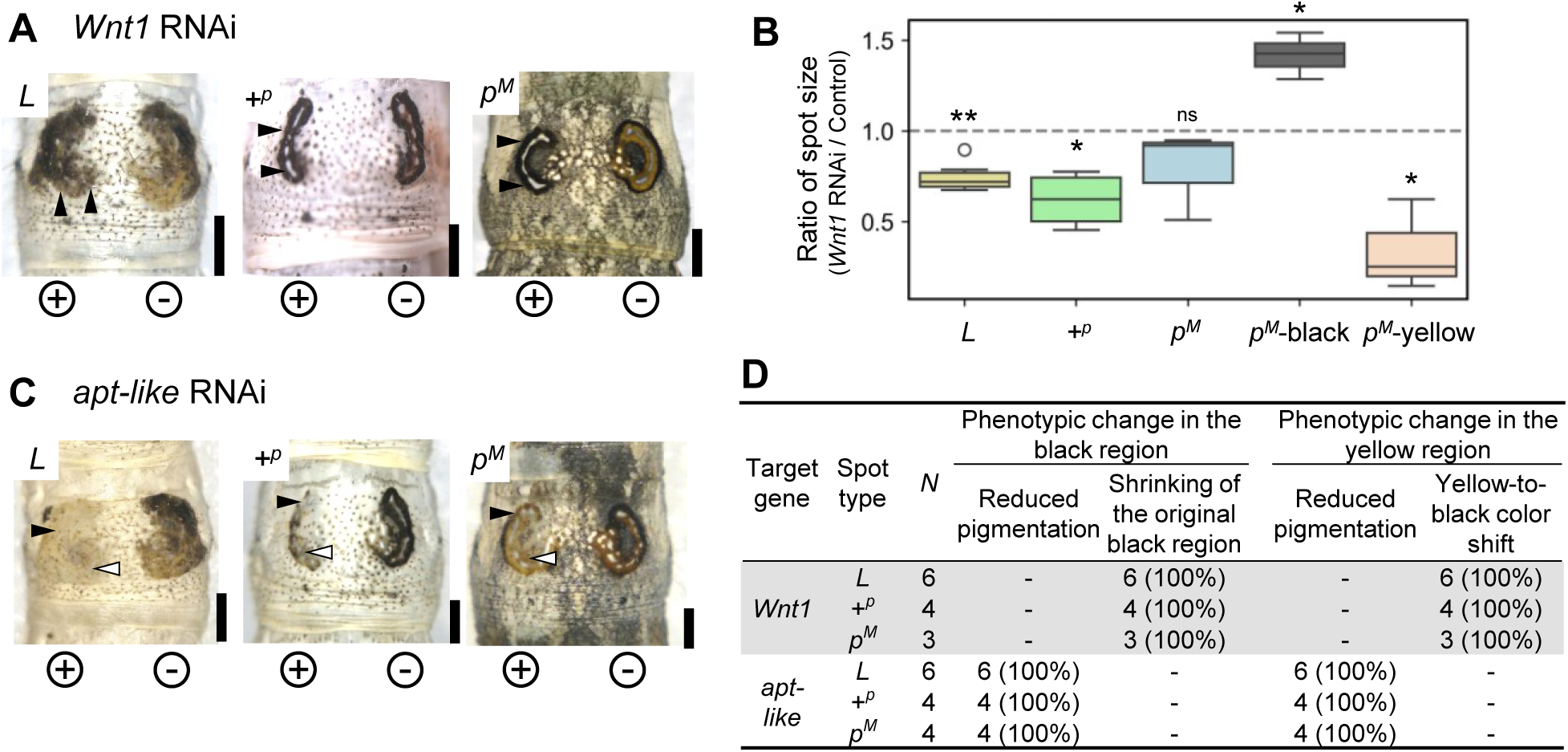
RNAi knockdown effect of *Wnt1* and *apt-like* on larval spots in the *L*-, +*^p^*-, and *p^M^*-hybrid strains. (**A**) *Wnt1* RNAi phenotypes in *L-*, +*^p^-*, and *p^M^*-hybrid spots. RNAi was applied to the left-side spots (plus electrode side) using *in vivo* electroporation. *Wnt1* knockdown resulted in partial or overall spot size reduction (arrowheads), primarily due to the shrinkage of the black-pigmented region. A yellow-to-black color shift was also observed in the inner area. (**B**) Spot size reduction quantification in *Wnt1* RNAi individuals. The spot areas on the untreated (right side) control were used as references for calculating the relative area of the RNAi-treated side. *L*, *L-*spots; +*^p^*, +*^p^-*spots; *p^M^*, *p^M^*-hybrid spots; *p^M^*-black, black region of the *p^M^*-hybrid spots; *p^M^*-yellow, yellow region of the *p^M^*-hybrid spots. Statistical significance was assessed using paired *t*-test: *P* < 0.05; *P* < 0.01; ns, not significant. (**C**) *apt-like* RNAi phenotypes. In contrast to *Wnt1* knockdown, *apt-like* silencing resulted in an almost complete loss of both black and yellow pigmentation (black and white arrowheads, respectively) in the left-side spots, without any spot size or inner color composition change. (**D**) Summary of *Wnt1* and *apt-like* RNAi phenotypes. The table summarizes pigmentation changes in the black and yellow regions upon gene knockdown. Scale bars: 1 mm.

Intriguingly, while the area of the black region expanded significantly (Fig. 5B, *p^M^*-black), the area of the yellow region significantly reduced (Fig. 5B, *p^M^*-yellow), implying that the yellow region may have been replaced by black pigmentation.

These results suggest that *Wnt1* expression levels influence both spot size and black and yellow melanin synthesis regulation. Notably, despite the reduced *Wnt1* expression in the *L-* spots, the primary observed effect was a spot size reduction rather than transformation into *+^p^-*like spots.

In contrast, *apt-like* knockdown in the *L-* and *+^p^-*spots reduced both black and yellow pigmentation without affecting spot size. I observed a similar effect in the *p^M^*-spots, confirming the role of *apt-like* in pigmentation regulation (Fig. 5C). Notably, while *Wnt1* knockdown did not impact the branch-like markings observed in *p^M^* hybrids, *apt-like* knockdown inhibited their formation. This contrast suggests that *Wnt1* primarily regulates spot formation, whereas *apt-like* might retain a broader role in controlling larval pigmentation patterns, including branch-like markings.

Figure 5D contains the compiled results, summarizing the effects of *Wnt1* and *apt-like* RNAi across *L-*, *+^p^-*, and *p^M^*-hybrid spots. This comparison highlights the differential roles of these genes in regulating both pigmentation and spot morphology.

To further validate the role of *apt-like* in black and yellow pigmentation, I performed a TALEN-mediated somatic knockout in the *p^M^*strain. I designed the TALEN target site within exon 6, encoding a leucine zipper-like motif essential for pigmentation induction (Yoda et al., 2014). Among 61 embryos injected with TALEN mRNA, three larvae hatched, and one exhibited white patches replacing black and yellow pigmentation in the *p^M^*-hybrid spots, along with disrupted branch-like body markings (Fig. S4). Sequencing of this mosaic larva revealed two distinct *apt-like* mutants, i.e., one with a 6-bp deletion (Δ6) and the other with a 16-bp deletion (Δ16). Notably, the Δ16 mutation likely introduced a frameshift, disrupting the coding region and generating a premature stop codon. These results collectively demonstrate that *apt-like* is essential for regulating black and yellow pigmentation in *p^M^*-hybrid spots and for the development of branch-like markings, but does not influence *p^M^*-hybrid spot size.

## 4. Discussion

In this study, I used qPCR and RNAi to assess the role of six core melanin synthesis genes in the black and yellow regions in *B. mori* larval spots. Similarly, I used qPCR to quantify *Wnt1* and *apt-like* expression and performed RNAi to investigate how these genes regulate both pigmentation types. My results suggest a potential gene regulatory network with *Wnt1* expression levels proving crucial for activating distinct melanin synthesis gene subsets via *Wnt1-apt-like* interactions, ultimately resulting in black and yellow pigmentation in the larval spots (Fig. 6).

**Fig. 6.**
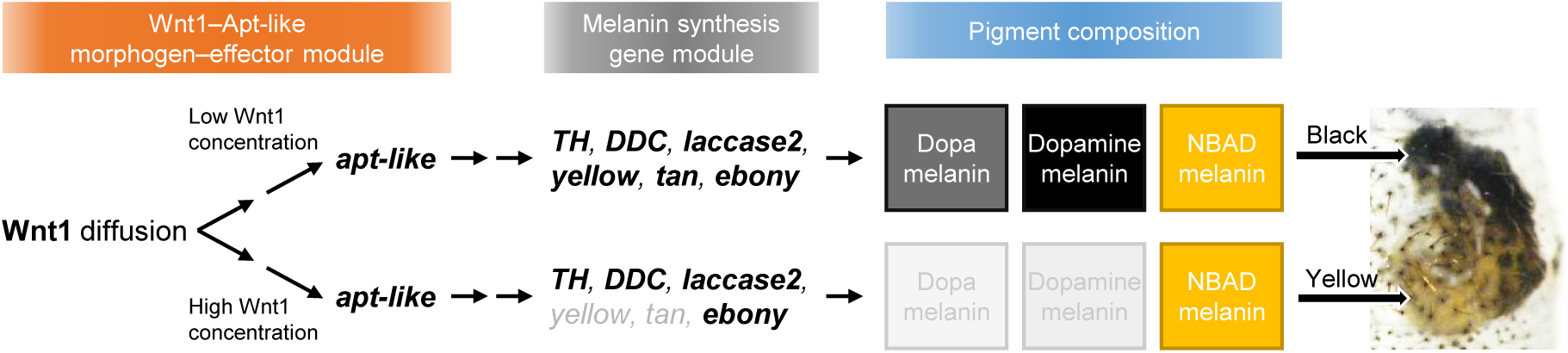
*Wnt1*-driven pigment differentiation through *apt-like* activation and melanin pathway modulation. Proposed gene regulatory model with *Wnt1* expression levels playing a central role in modulating distinct melanin synthesis gene subsets via the Wnt1–Apt-like axis, ultimately determining black and yellow pigmentation in the *L*-spots. Both low and high Wnt1 levels induce *apt-like* expression. However, they differentially activate downstream melanin synthesis genes (highlighted in bold), leading to the production of distinct melanin types. In the black regions, both dopa/dopamine-melanin and NBAD-melanin are synthesized, whereas the yellow regions are predominated by NBAD-melanin. The intermediate regulatory steps connecting *Wnt1* to *apt-like*, as well as those linking the Wnt1–Apt-like module to the specific activation of melanin synthesis genes, remain unclear.

My discoveries demonstrate that black and yellow pigmentation in *B. mori* larval spots is regulated by distinct yet overlapping melanin synthesis gene sets. qPCR and RNAi experiments revealed that *TH*, *DDC*, *laccase2*, *ebony*, *yellow*, and *tan* contribute to the formation of the black region, whereas *TH*, *DDC*, *laccase2*, and *ebony* are essential for the yellow region (Figs. 2 and 3). These results indicate that black pigmentation involves both dopa/dopamine- and NBAD-melanin pathways, while yellow predominantly requires only the latter. The modular use of melanin synthesis genes I identified in this study aligns well with genetic architectures previously observed in other insects. For example, in *Papilio xuthus*, black and reddish-brown pigmentations in the larval eyespots are similarly regulated by partially overlapping melanin-related gene sets during the final larval molt (Futahashi et al., 2010). Modular melanin pathway regulation has also been demonstrated through functional analyses in nymphalid butterflies such as *Vanessa cardui* and *Bicyclus anynana* (Zhang et al., 2017; Matsuoka and Monteiro, 2018) as well as in hemimetabolous insects including *Oncopeltus fasciatus* and *Zootermopsis nevadensis* (Liu et al., 2016; Masuoka and Maekawa, 2016). However, unlike previously studied species, my results in *B. mori* reveal that *ebony* contributes both to black and yellow pigmentation, raising the possibility that darkness- or yellowishness-related differences depend on the relative dopa/dopamine-melanin-to-NBAD-melanin proportion synthesized within each region. In addition, the RNAi phenotypes suggest that melanin composition differences might influence the boundary sharpness between black and yellow areas (Fig. 3). Taken together, these results highlight how differential gene usage within conserved melanin synthesis pathways generates diverse pigmentation outcomes in insects.

Consistent with its proposed role in defining pigmentation domains (Yamaguchi et al., 2013), *Wnt1* expression is spatially restricted to presumptive spot regions in *B. mori* (Fig. 4A). *Wnt1* RNAi resulted in reduced spot size and yellow-to-black color shift (Figs. 5A and B), suggesting that *Wnt1* influences both spatial prepatterning and pigmentation composition.

This *Wnt1* role in the silkworm mirrors its function in butterfly wings, where Wnt signaling establishes spatial pigmentation patterns, such as that of eyespots (Monteiro et al., 2006; Monteiro, 2015; Özsu et al., 2017; Mazo-Vargas et al., 2017; Banerjee et al., 2023). Similar to butterflies, Wnt1 appears to act dose-dependently in silkworms, regulating both the pigmented domain size and produced pigment type. This supports the idea that Wnt1 functions as a prepatterning module, defining pigmentation domains through a concentration-dependent mechanism. As a morphogen, Wnt1 could form a localized gradient originating from a central source, such as the bulge structure proposed by Yamaguchi et al. (2013). In this model, cells near the source are exposed to higher Wnt1 levels, activating gene subsets that initiate and expand the spot, while cells farther away respond to lower Wnt1 levels, thereby contributing to the outer boundaries of the spot.

Beyond regulating spot size, Wnt1 concentration also appears to influence pigmentation type. Higher Wnt1 levels activate downstream genes associated with yellowish pigmentation, while lower levels activate a partially overlapping set of more sensitive genes involved in darker pigmentation. This result is broadly consistent with models, in which Wnt functions as a morphogen that diffuses from source cells and establishes patterning gradients in wing tissues (Monteiro et al., 2006; Werner et al., 2010; Martin et al., 2012). In butterfly eyespot formation (e.g., in the case of *B. anynana*) a “gradient model” has been proposed, in which morphogen concentration thresholds define distinct pigment zones, resulting in a sharp transition between differently colored rings (Brunetti et al., 2001; Allen et al., 2008; Özsu et al., 2017). In contrast, my findings in *Bombyx L*-spots suggest a more gradual, concentration-dependent response to Wnt1. In this model, rather than being separated by a strict threshold, black and yellow pigmentation emerge continuously along a Wnt1 gradient. The observed yellow-to-black color shift upon *Wnt1* knockdown in the *L*-spots supports this idea and likely reflects a shift in the balance between melanin types. High Wnt1 levels may promote NBAD-melanin synthesis by upregulating *TH*, *DDC*, *laccase2*, and *ebony*, whereas lower levels may promote both NBAD-melanin and dopa/dopamine-melanin production through the upregulation of partially overlapping genes (*TH*, *DDC*, *laccase2*, *ebony*) and additional genes such as *yellow* and *tan*. This mixed melanin composition in the black region results in a darker pigmentation that is visually distinct from the yellow region, despite the shared involvement of NBAD-melanin (Fig. 6). Such dosage-sensitive regulation is consistent with Wnt1 functioning as a morphogen that fine-tunes pigmentation patterns through the graded control of melanin synthesis pathways.

However, the extent of overall spot identity determination by Wnt1 concentration alone remains unclear. While my knockdown experiments indicated that reducing *Wnt1* levels in *L*-spots caused a noticeable spot size reduction and altered pigmentation, it did not lead to *L*-spot transformation into crescent-shaped +*^p^*-spots, as might have been expected. If high Wnt1 expression were solely responsible for *L*-spot identity specification, then its reduction should at least partially shift the *L*-spot toward a +*^p^*-like state. Instead, the partial phenotypic change implies that *L-*spot fate is maintained by additional regulatory mechanisms beyond Wnt1 signaling during the larval period. A possible explanation is that the difference in *L-* spots versus +*^p^-*spots identity might be established during early embryogenesis, wherein *Wnt1* expression (and possibly that of other factors) act during a critical time window to define prospective *L-*spot source cell fates. Aruga et al. (1954) referred to these embryonic precursors as an “invisible anlage,” underscoring that they are not visibly distinct at early stages but already primed for larval spot formation. Once these cells adopt an *L-*spot “ground state,” reducing *Wnt1* levels at later stages might not suffice to switch them to a +*^p^-* spot fate. This scenario suggests an epigenetic “lock-in” of the *L-*spot identity triggered by *Wnt1*- and other regulatory input-involving early developmental events.

The regulatory mechanism by which *apt-like* contributes to both black and yellow pigmentation remains unclear. *apt-like* expression in the *+^p^* allele is suggestibly controlled by a *cis*-regulatory element located within a 33-kb region of the *apt-like* locus, presumably mediating its responsiveness to *Wnt1* signaling (Yoda et al., 2014). This concept is supported by observations that *Wnt1* expression correlates with *apt-like* activation, and that ectopic *Wnt1* expression induces *apt-like*, whereas ectopic *apt-like* expression does not induce *Wnt1* (Yoda et al., 2014). These observations support a model in which *Wnt1* functions as an upstream *apt-like* regulator. In this study, I observed that *apt-like* expression levels differed between pigment regions, with higher expressions in the black than in the yellow region (Fig. 4B). Although further regulatory factors might also be involved, this expression difference likely reflects variation in Wnt1 signaling activity between the black and yellow regions.

To further reveal the *Wnt1–apt-like* regulatory network, investigating other potential contributor signaling or transcriptional pathways to this system would be important. One such candidate is the Notch signaling pathway. *Notch* is reportedly required for pigmentation, as its RNAi knockdown results in the loss of both black and yellow pigments in the *L-*spots (Jin et al., 2020). However, the regulatory relationship between *Notch* and the *Wnt1–apt-like* axis remains unresolved. Future studies incorporating comprehensive transcriptomic approaches (e.g., RNA-sequencing across multiple developmental stages and pigment regions, with and without knockdowns of these key genes as well as ATAC-seq to identify open chromatin regions) will be essential for unravelling the broader gene regulatory network underlying larval spot formation.

## Acknowledgements

The experimental work for this study was primarily conducted at the Department of Integrated Biosciences, Graduate School of Frontier Sciences, The University of Tokyo (Kashiwa, Japan), where I was affiliated from 2015 to 2020. I am grateful to Drs. Haruhiko Fujiwara, Tetsuya Kojima, Yusuke Kondo, and Pierre Galipot for their valuable support during that period. Silkworm strains used in this study were assisted by the National Bio-Resource Project (NBRP) of the MEXT, Japan.

---

**Supplementary Fig. 1.**
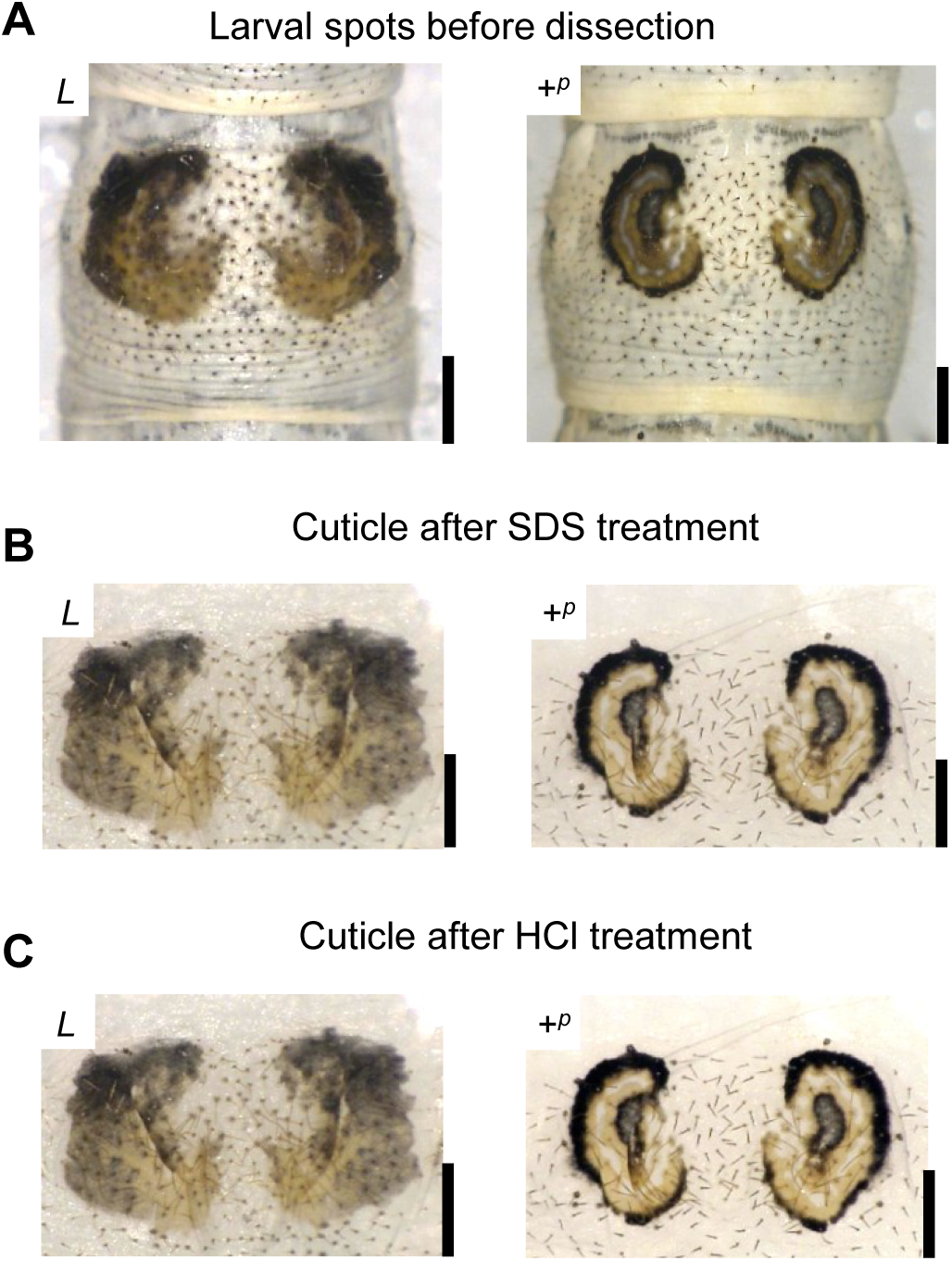
Estimation of pigment composition in larval spots using chemical treatments. (**A**) Natural pigmentation patterns of larval spots in *L* and +*^p^* phenotypes prior to chemical treatment. (**B**) After removal of the epidermal cells by SDS treatment, both black and yellow pigmentation remain visible in the cuticle. (**C**) HCl treatment of SDS-treated cuticle specimens does not eliminate the black or yellow pigments, indicating that both are composed of highly polymerized, acid-resistant melanin. Scale bars, 1 mm.

**Supplementary Fig. 2.**
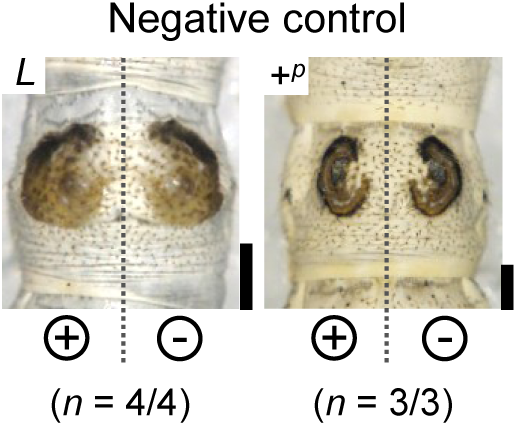
Negative control for *in vivo* electroporation-mediated RNAi. No phenotypic changes were observed in *L-* and *+^p^*-spots following the negative control treatment. Scale bars, 1 mm.

**Supplementary Fig. 3.**
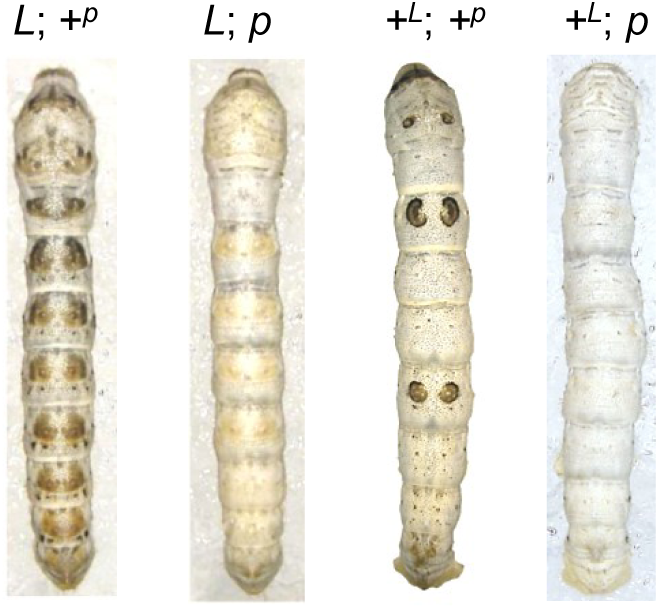
Allelic influence of the *p* locus on *L*- and +*^p^*-spot formation. Allelic variation at the *p* locus modulates *L*-spot formation. *L*-spots can be observed when the +*^p^* allele is present, either in the homozygous (+*^p^*/+*^p^*) or heterozygous (+*^p^*/*p*) state (*L*; +*^p^*). In contrast, *L*-spots are absent in homozygous *p*/*p* individuals (*L*; *p*), indicating that *L*-spot expression requires at least one functional +*^p^* allele. In the absence of the *L* mutation, individuals carrying the +*^p^* allele (+*^L^*; +*^p^*) develop +*^p^*-spots, whereas those lacking it (+*^L^*; *p*) do not form +*^p^*-spots.

**Supplementary Fig. 4.**
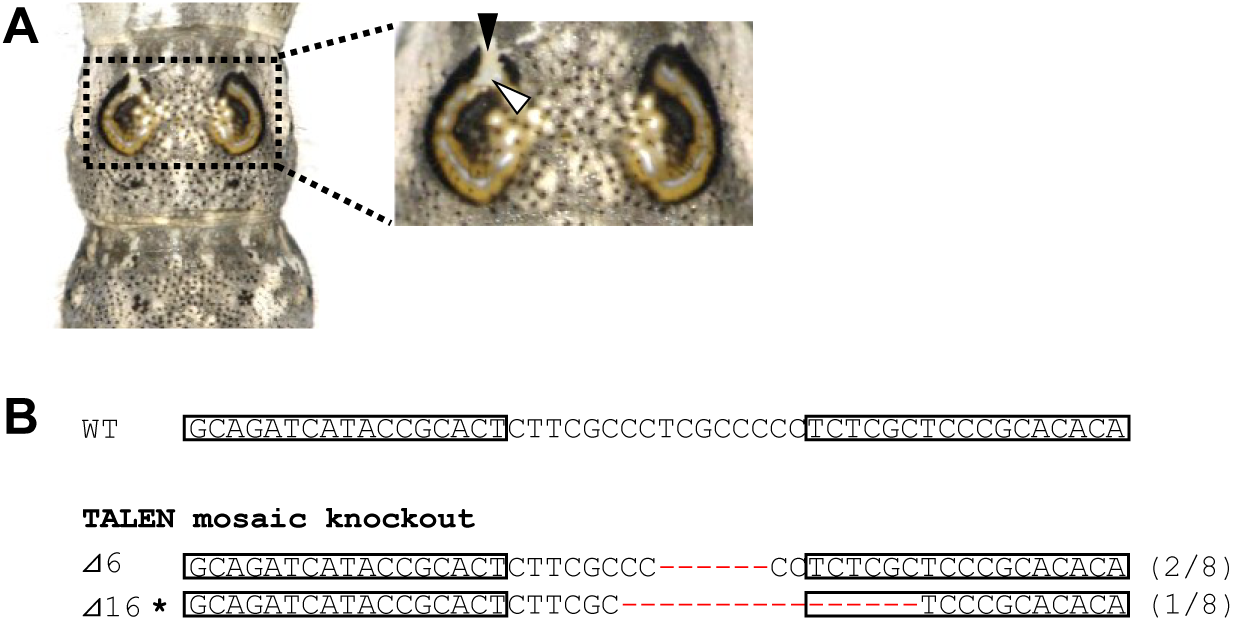
TALEN-mediated somatic knockout of *apt-like* in a *p^M^* introgression strain. (**A**) Mosaic loss of pigmentation was observed in a *p^M^*-spot following TALEN-mediated knockout of *apt-like*, including partial loss of black pigmentation (black arrowhead) and yellow pigmentation (white arrowhead). (**B**) TALEN-induced mutations resulted in deletions surrounding the TALEN-binding sites (indicated by boxes). The wild-type (WT) sequence is shown above two representative mutant sequences, with deletions indicated by red dashes. Asterisks denote a frameshift mutation. The number of clones carrying each mutation is shown to the right of each sequence. *Note: The strain used here is an introgression line in which the p^M^ allele derived from Bombyx mandarina was backcrossed into a B. mori genetic background; the B. mandarina genome is retained only on chromosome 2*.

**Supplementary Table 1.**
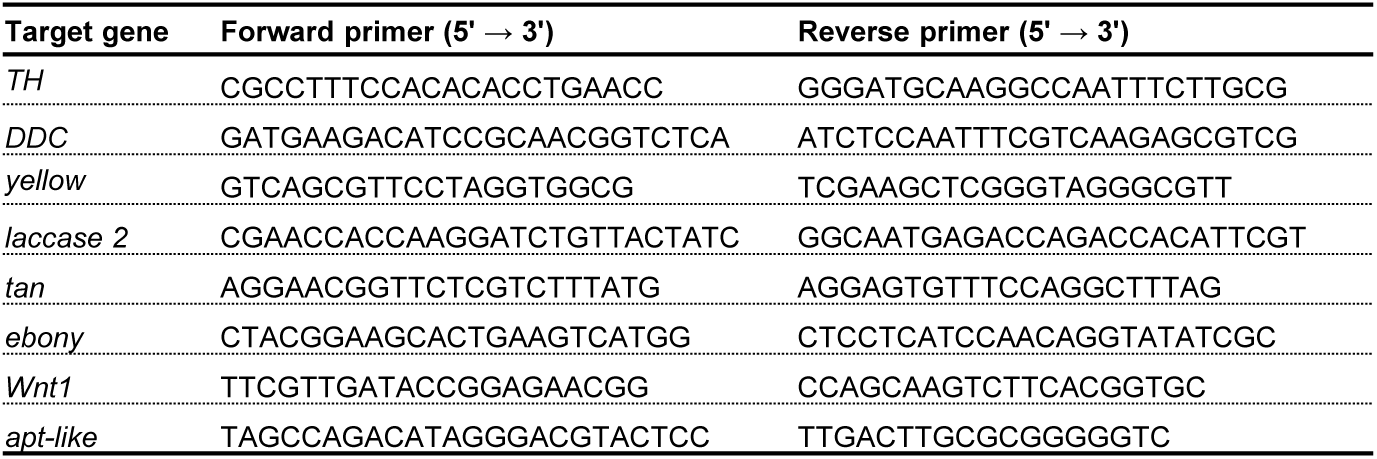
Primers used for qPCR.

**Supplementary Table 2.**
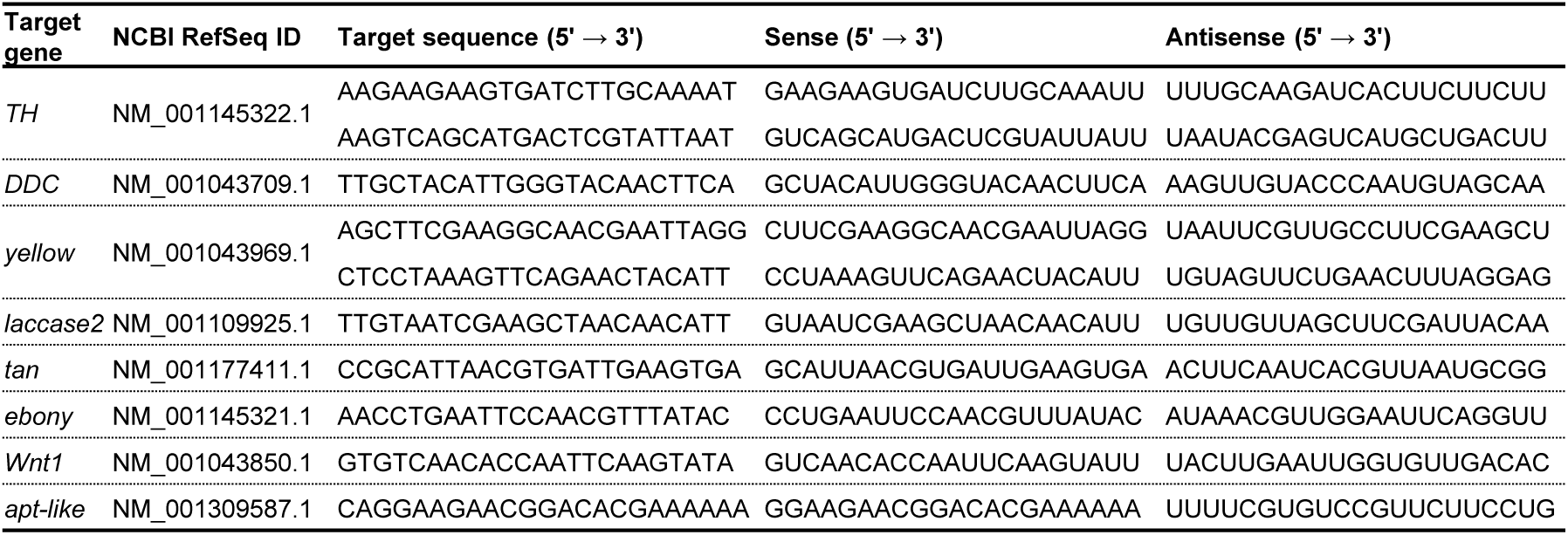
siRNAs used for RNAi.

